# G2S3: a gene graph-based imputation method for single-cell RNA sequencing data

**DOI:** 10.1101/2020.04.01.020586

**Authors:** Weimiao Wu, Qile Dai, Yunqing Liu, Xiting Yan, Zuoheng Wang

## Abstract

Single-cell RNA sequencing provides an opportunity to study gene expression at single-cell resolution. However, prevalent dropout events result in high data sparsity and noise that may obscure downstream analyses. We propose a novel method, G2S3, that imputes dropouts by borrowing information from adjacent genes in a sparse gene graph learned from gene expression profiles across cells. We applied G2S3 and other existing methods to seven single-cell datasets to compare their performance. Our results demonstrated that G2S3 is superior in recovering true expression levels, identifying cell subtypes, improving differential expression analyses, and recovering gene regulatory relationships, especially for mildly expressed genes.

## Background

Singe-cell RNA sequencing (scRNA-seq) has emerged as a state-of-the-art technique for transcriptome analysis. Compared to bulk RNA-seq that measures the average gene expression profile of a mixed cell population, scRNA-seq measures cellular level expression for each gene and thus describes cell-to-cell stochasticity in gene expression. Applications of this technology in humans have revealed rare or novel cell types [1–3], cell population compositional changes [4], and cell-type specific transcriptomic changes [3,5] that are associated with diseases. These findings have great potential to promote our understanding of cell function, disease pathogenesis, and treatment response for more precise therapeutic development [6,7]. However, analysis of single-cell transcriptomic data can be challenging due to low library size, high noise level, and prevalent dropout events [8]. Particularly, dropouts lead to an excessive number of zeros or close to zero values in the data, especially for genes with low or moderate expression. These inaccurately measured gene expression levels may obscure downstream quantitative analyses such as cell clustering and differential expression analyses [6].

In the past few years, several imputation methods have been developed to recover dropout events in single-cell transcriptomic data. A group of methods, including MAGIC [9], scImpute [10], drImpute [11], and VIPER [12], assess between cell similarity and impute dropouts in each cell using its similar cells. Specifically, MAGIC constructs an affinity matrix of cells and aggregates gene expression across similar cells via data diffusion to impute gene expression for each cell [9]. scImpute infers dropout events based on the dropout probability estimated from a Gamma-Gaussian mixture model and only imputes these events by borrowing information from similar cells within cell clusters detected by spectral clustering [10]. drImpute identifies similar cells through K-means clustering and performs imputation via averaging the expression levels of cells within the same cluster [11]. While these imputation methods improved the quality of single-cell transcriptomic data to some extent, they were found to eliminate natural cell-to-cell stochasticity which is an important piece of information available in scRNA-seq data compared to bulk RNA-seq data [12]. To overcome this limitation, VIPER considers a sparse set of neighborhood cells for imputation to preserve variation in gene expression across cells [12]. In general, imputation methods that borrow information across similar cells tend to intensify subject variation in scRNA-seq datasets with multiple subjects when cells from the same subject are more similar than those from different subjects. To address this issue, SAVER borrows information across similar genes instead of cells to impute gene expression using a penalized regression model [13]. In addition, machine learning-based methods such as autoImpute [14], DAC [15], and deepImpute [16], use deep neural network to impute dropout events. While computationally more efficient, these methods were found to generate false-positive results in differential expression analyses [17].

In this article, we develop G2S3, a sparse and smooth signal of gene graph-based method that imputes dropout events in single-cell transcriptomic data by borrowing information across similar genes. G2S3 learns a sparse graph representation of gene-gene relationships from scRNA-seq data, in which each node represents a gene and is associated with a vector of expression levels in all cells that can be viewed as a signal on the graph. The graph is then optimized under the assumption that signals change smoothly between connected genes. Based on this graph, a transition matrix for a random walk is constructed so that the transition probabilities between genes with similar expression levels across cells are higher. A random walk on this graph imputes the expression level of a gene using the weighted average of expression levels from itself and adjacent genes in the graph. In this way, G2S3, like SAVER, makes use of gene-gene relationships to recover the true expression levels. However, unlike SAVER which uses a penalized regression model for imputation, G2S3 optimizes the gene graph structure using graph signal processing that captures nonlinear correlations among genes and is robust to outliers in the data. The computational complexity of the G2S3 algorithm is a polynomial of the total number of genes in the graph, so it is computationally efficient, especially for large scRNA-seq datasets with hundreds of thousands of cells.

## Results

### Datasets and analyses overview

We evaluated and compared the performance of G2S3 and five existing imputation methods, SAVER, MAGIC, scImpute, VIPER, and DCA, in terms of (1) expression data recovery; (2) cell subtype separation; (3) differential gene identification; and (4) gene regulatory relationship recovery. We applied these imputation methods to seven scRNA-seq datasets that can be classified into four categories accordingly. The first category includes three unique molecular identifier (UMI)-based datasets for the down-sampling analysis: the Reyfman dataset from human lung tissue [18], the peripheral blood mononuclear cell (PBMC) dataset from human peripheral blood [19], and the Zeisel dataset from mouse cortex and hippocampus [20]. Down-sampling was applied to these datasets to assess the method performance in recovering true expression levels. The second category of datasets was used to evaluate the method performance in separating different cell types. It includes the Chu dataset of human embryonic stem (ES) cell-derived lineage-specific progenitors from seven known cell subtypes [21], and the Petropoulos dataset of cells from human preimplantation embryos collected on different embryonic days [22]. The third category was chosen to evaluate the method performance in improving the identification of differentially expressed genes. It includes the Segerstolpe dataset of pancreatic islets from type II diabetes (T2D) patients and healthy controls [23]. The last category includes the dataset from Paul et al. [24] that measured the well-known transcriptional regulators of myeloid progenitor populations. With this dataset, we evaluated the method performance in restoring gene regulatory relationships among key regulators. Table 1 summarizes the main features of the seven datasets. A more detailed description of these datasets is provided in the “Real datasets” section.

**Table 1.** Detailed information on the seven scRNA-seq datasets used to compare the performance of imputation methods

| Experiment Category | Dataset | # Cells | Sample Type | Organism | Technique | UMI | Accession |
| --- | --- | --- | --- | --- | --- | --- | --- |
| Expression data recovery | Reyfman [18] | 5,437 | Lung tissue | Homo Sapiens | Drop-seq | Yes | GEO (GSE122960) |
|  | PBMC [19] | 7,865 | Peripheral blood mononuclear cells | Homo Sapiens | Drop-seq | Yes | 10x Genomics* |
|  | Zeisel [20] | 3,005 | Brain tissue | Mus Musculus | Drop-seq | Yes | Zeisel et al. |
| Cell subtype separation | Chu [21] | 1,018 | Embryonic stem cells | Homo Sapiens | Fluidigm C1 | No | GEO (GSE75748) |
|  | Petropoulos [22] | 1,529 | Preimplantation embryos | Homo Sapiens | Smart-seq2 | No | Petropoulos et al. |
| Differential gene identification | Segerstolpe [23] | 3,514 | Whole islets | Homo Sapiens | Smart-seq2 | No | Segerstolpe et al. |
| Gene regulatory relationship recovery | Paul [24] | 2,730 | Bone marrow myeloid progenitor | Mus Musculus | MARS-seq | Yes | Paul et al. |

\* URL to access the dataset: https://support.10xgenomics.com/single-cell-gene-expression/datasets

### Expression data recovery in down-sampled datasets

Using the three down-sampled scRNA-seq datasets (Reyfman, PBMC, and Zeisel), we assessed the performance of the six imputation methods in recovering true expression levels. Figure 1 shows the gene-wise Pearson correlation and cell-wise Spearman correlation between the imputed and reference data using each dataset. The correlation between the observed data without imputation and reference data was set as a benchmark. In all datasets, G2S3 consistently achieved the highest correlation with the reference data at both gene and cell levels, and SAVER had slightly worse performance. VIPER performed well in the Reyfman and PBMC datasets but not in the Zeisel dataset based on gene-wise correlation, although the cell-wise correlations were much lower than G2S3 and SAVER in all datasets. MAGIC, scImpute, and DCA did not have comparable performance, especially based on gene-wise correlation. Since genes with higher expression tend to have a lower dropout rate, they are usually easier to impute but have less imputation need than those with lower expression [8]. To demonstrate the impact of expression level on the method performance, we stratified genes into three subsets based on the proportion of cells expressing them in the reference data: widely expressed (>80%), mildly expressed (30%-80%), and rarely expressed (<30%). Figure S1 shows the gene-wise and cell-wise correlations in each gene stratum. We can see that G2S3 improved both the gene-wise and cell-wise correlations compared to the observed data for the widely and mildly expressed genes. Moreover, G2S3 achieved the most superior recovery accuracy than the other methods for the widely and mildly expressed genes, although SAVER had comparable accuracy for the widely expressed genes, suggesting the advantage of borrowing information from similar genes over from similar cells. For the rarely expressed genes, all imputation methods did not improve the correlations compared to the observed data at both gene and cell levels, suggesting that there is insufficient information on these genes to be successfully imputed. Overall, G2S3 provided the most accurate recovery of true expression levels.

**Figure 1.**
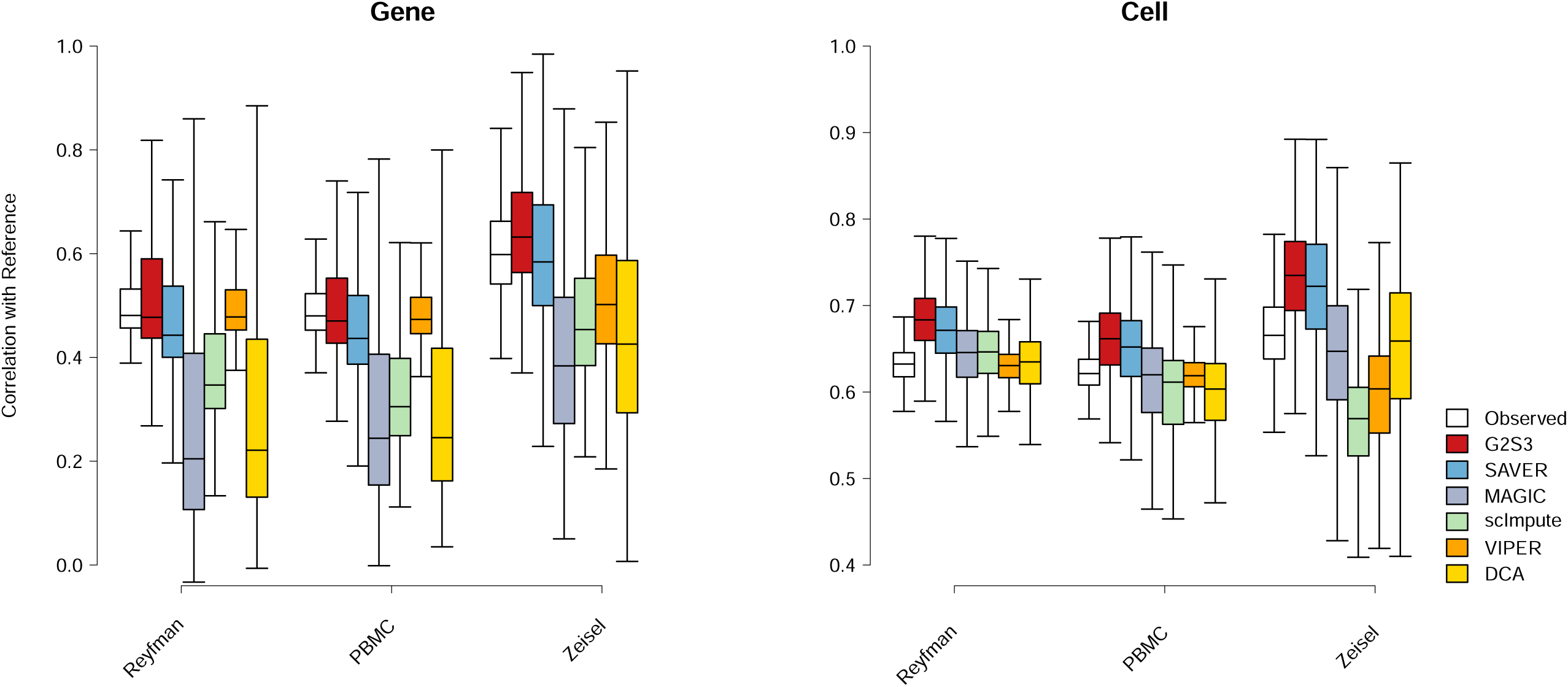
Performance of imputation methods measured by correlation with reference data in three down-sampled datasets, on the gene level (left) and cell level (right). Box plots show the median (center line), interquartile range (hinges), and 1.5 times the interquartile (whiskers), outlier data beyond this range are not shown.

### Restoration of cell subtype separation

The second category of datasets was used to assess the performance of imputation methods in restoring the separation between different cell types. In the Chu dataset, there were 7 cell types including two undifferentiated human ES cell lines (H1 and H9), human foreskin fibroblasts (HF), neuronal progenitor cells (NP), definitive endoderm cells (DE), endothelial cells (EC), and trophoblast-like cells (TB). To quantify the performance in separating these cell subtypes, we calculated the ratio of the average inter-subtype distance to the average intra-subtype distance using the top *K* principal components (PCs) of the data before and after imputation, for *K* = 1, …, 10. We also calculated the silhouette coefficient that measures how similar cells are to cells from the same cell type compared to other cell types. Note that SAVER was unable to finish imputation within 24 hr on this dataset and thus was not shown. In Figure 2, G2S3 had the highest inter/intra-subtype distance ratio, and VIPER was the second best. Both methods performed better than the raw unimputed data, while MAGIC, scImpute, and DCA performed worse than the raw data. G2S3 also achieved the highest silhouette coefficient among all imputation methods for all *K*. These results suggest that G2S3 greatly improved the separation between different cell types and achieved the best performance than the other imputation methods.

**Figure 2.**
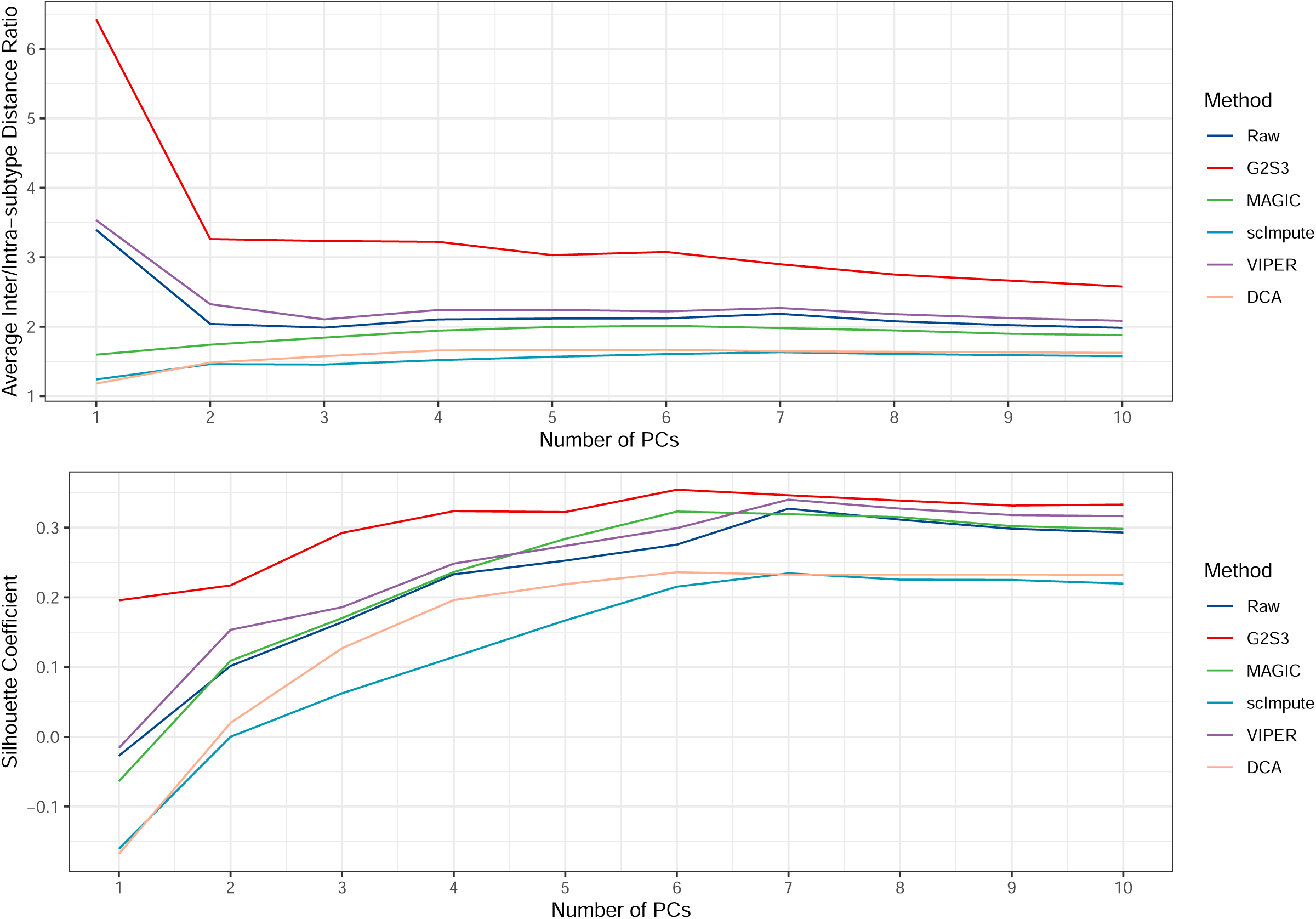
Average inter/intra-subtype distance ratio (top) and silhouette coefficient (bottom) to demonstrate cell subtype separation using the top *K* principal components of the raw unimputed and imputed data by each method in the Chu dataset.

To demonstrate the comparison using cell clustering results, we generated PC plots in which cells were colored to represent the seven cell subtypes in the original dataset. Figures 3 and S2 showed that the imputed data by G2S3 generated better separation of all cell subtypes except H1 and H9 cells than the raw unimputed data. Given that both H1 and H9 are undifferentiated human ES cell lines, it is expected that separating them is more difficult due to the relative homogeneity of human ES cells compared to the progenitors. In contrast, the other imputation methods did not have comparable improvement or even reduced the separation of different cell types. Specifically, DE cells were mixed with EC and TB cells in the raw dataset and were not separated from the other cell subtypes by all methods except G2S3. MAGIC was only able to separate EC, H1 and HF cells from each other and the rest of the cell subtypes. scImpute tended to mix different cell types into one cluster. VIPER increased the within-subtype variation and resulted in artificially more heterogeneous cell-type clusters. DCA was only able to separate H1/H9 and HF cells from the rest.

**Figure 3.**
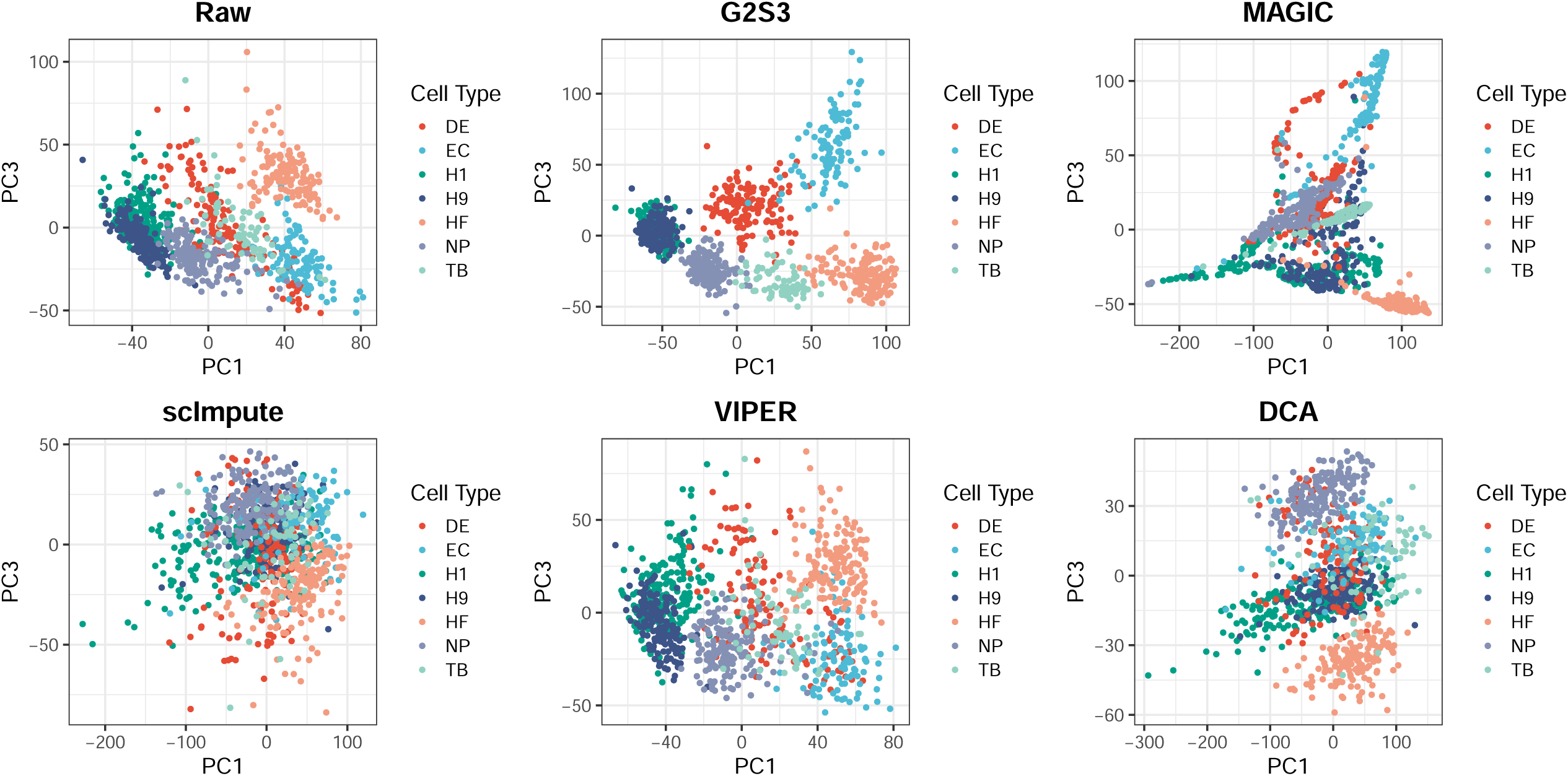
PC plots (PC1 vs. PC3) of the raw unimputed and imputed data by each method in the Chu dataset. Cells are colored by the cell subtype labels.

Figure S3 demonstrated the expression of two cell subtype marker genes *GATA*6, a marker gene of DE cells, and *NANOG*, a marker gene of H1/H9 cells [21] across all cells. We can see that G2S3 provided the best separation between H1/H9 cells, DE cells and other cell subtypes. Specifically, while neither the raw data nor the other imputation methods showed a clear separation between DE and NP cells, G2S3 successfully separated these two cell subtypes both from each other and from the other cell subtypes. In addition, the imputed data by VIPER still had a large proportion of dropout events. DCA separated H1/H9 cell lines from the rest of the progenitor cells but had EC and TB cells marginally overlapped. These results suggest that G2S3 had the best performance in restoring the separation of different cell types, preserving biological meaningful variations, and reducing technical noises.

In the Petropoulos dataset, cells from different embryonic days (E3-E7) were considered as different cell types. We used t-distributed stochastic neighbor embedding (t-SNE) visualization to evaluate the separation between cell types. Given the performance advantage of G2S3 and MAGIC over the other methods in the Chu dataset, we only compared the performance of G2S3 and MAGIC in this dataset. The results showed that cells from stages E6 and E7 were mixed and inseparable via cell clustering using the Louvain algorithm [25,26] in the raw data (Figure 4A). After imputation by G2S3, these two stages of cells were separated as two clusters and the segregation of cells from different embryonic days was also generally improved (Figure 4B). In contrast, the imputed data by MAGIC resulted in tight clusters of cells that correspond to different subjects rather than biologically meaningful cell types (Figure 4C). This is likely because MAGIC powers up the Markov matrix built on the cell-to-cell affinity matrix, rendering cells within each subject more similar to each other.

**Figure 4.**
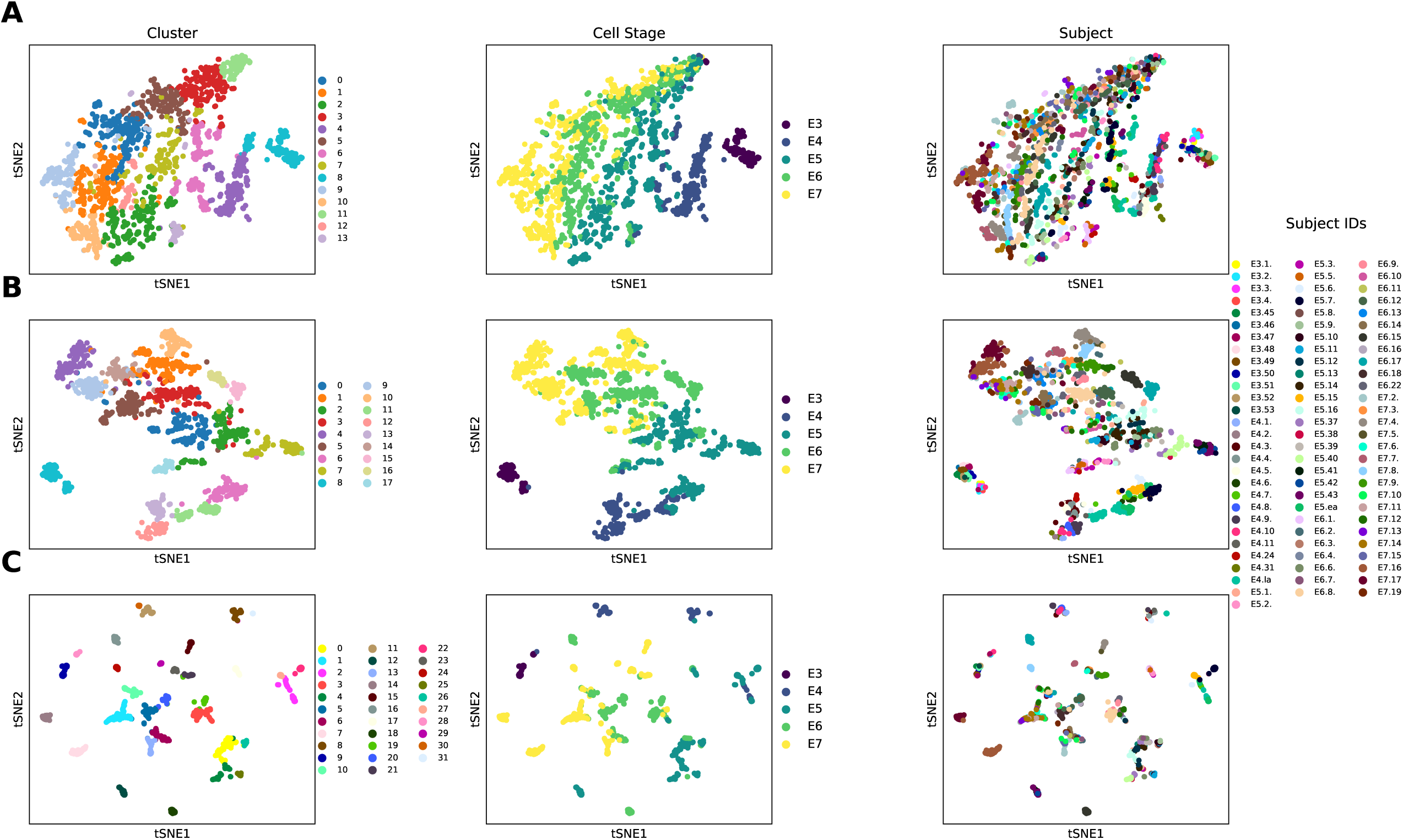
Cell clustering and t-SNE visualization of the Petropoulos dataset on human embryo stem cells from different embryonic days. **A**) t-SNE visualization of the raw data, **B**) t-SNE visualization of the imputed data by G2S3, and **C**) t-SNE visualization of the imputed data by MAGIC. In each panel, cells were colored by cluster ID detected via the Louvain clustering algorithm (left), cell embryonic day (middle), and subject ID (right).

### Improvement in differential expression analysis

One common analytical task for scRNA-seq studies is to identify genes differentially expressed between cells from two groups of subjects or two cell types. In this section, we evaluated and compared the improvement in downstream differential expression analysis before and after imputation by the six methods using the Segerstolpe dataset. Besides the scRNA-seq data, this dataset also provides bulk RNA-seq data on the same seven samples. We performed differential expression analysis between four T2D patients and three healthy donors using both scRNA-seq and bulk RNA-seq data. The differentially expressed genes identified from the bulk RNA-seq data were treated as ground truth. We assessed the predictive power of the scRNA-seq data imputed by different methods on the ground truth using receiver operating characteristic (ROC) curves. Genes were stratified into highly expressed (expressed in more than 30% cells, n=7,459) and lowly expressed groups (expressed in less than 30% cells, n=1,551) to examine the impact of expression level on the performance. Note that VIPER was unable to finish imputation within 24 hr and thus was not shown. Figure 5 demonstrated that for highly expressed genes all imputation methods did not improve the differential analysis results except MAGIC. One possible explanation, as was noticed in previous studies [10,12], is that MAGIC tends to overly smooth the data, making the imputed data artificially resemble the bulk RNA-seq data. For lowly expressed genes, all imputation methods improved the differential analysis results over the raw data. Among all methods, G2S3 achieved the highest area under the curve (AUC) using both the t-test and Wilcoxon test. SAVER obtained high AUC on the lowly expressed genes only in the Wilcoxon test results. DCA had the lowest AUC in both tests. MAGIC failed to detect any differentially expressed genes in the lowly expressed group using the scRNA-seq data. In summary, G2S3 achieved the best improvement in differential expression analysis, especially for lowly expressed genes.

**Figure 5.**
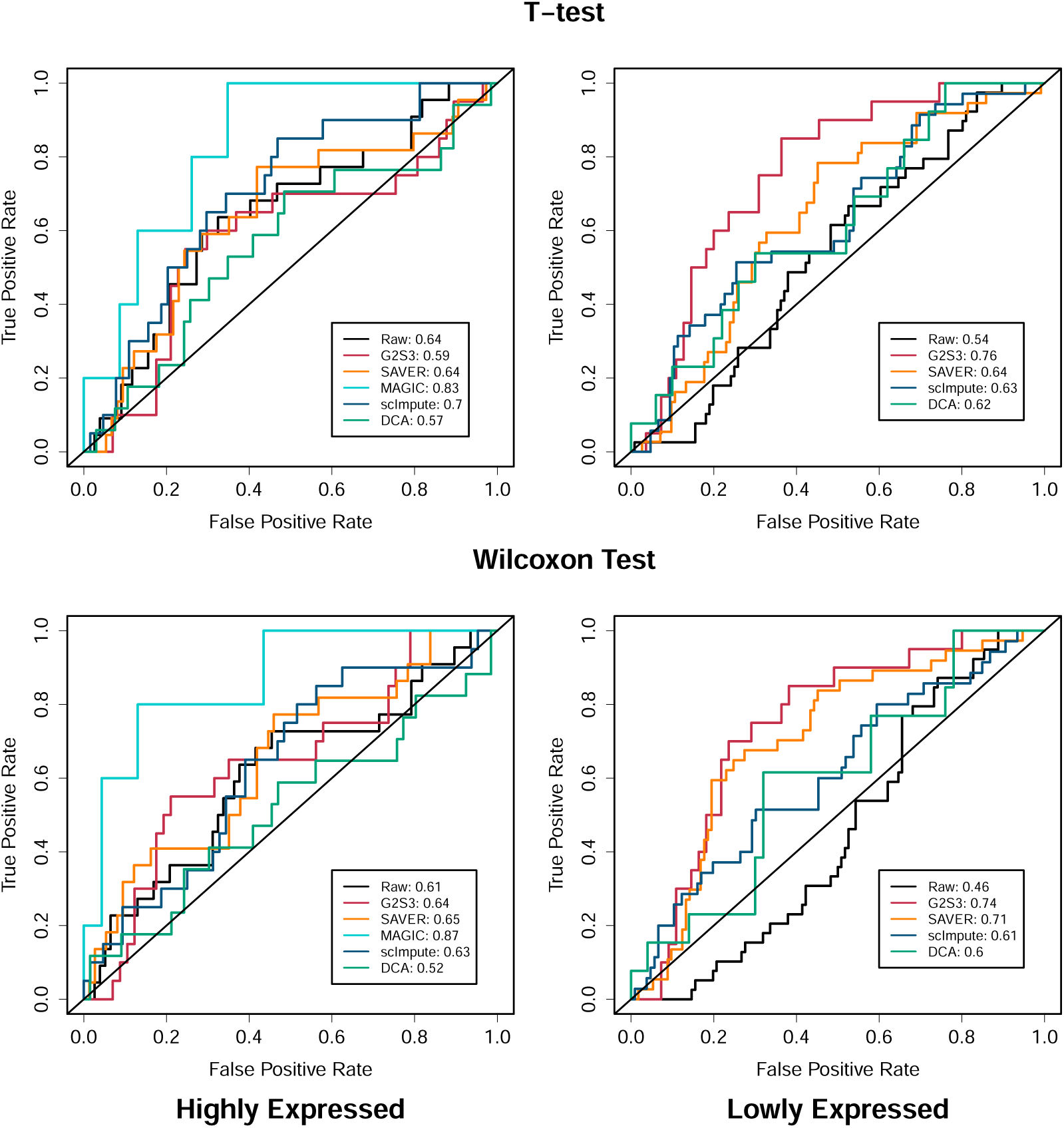
ROC curves of differential analysis results of the scRNA-seq data predicting differentially expressed genes identified from the bulk RNA-seq data in the Segerstolpe dataset. Genes are stratified into two groups: highly expressed (left) and lowly expressed (right).

### Gene regulatory relationship recovery

To compare the method performance in recovering gene regulatory relationships, we examined the pairwise correlation between well-known transcription factors in the development of blood cells using the Paul dataset. The pairwise correlations between key regulators of the transcriptional differentiation of megakaryocyte/erythrocyte progenitors and granulocyte/macrophage progenitors in the raw data and the imputed data by each method were used for performance evaluation. Based on previous studies [27–29], inhibitory and activatory gene pairs were defined, among which inhibitory pairs were expected to have negative correlation while activatory pairs were expected to have positive correlation. The mutually inhibitory pairs of genes include *Fli*1 vs. *Klf*1, *Egr*1 vs. *Gfi*1, *Cepbpa* vs. *Gata*1, and *Sfpi*1 vs. *Gata*1; and the mutually activatory pairs include *Sfpi*1 vs. *Cebpa, Zfpm*1 vs. *Gata*1, *Klf*1 vs. *Gata*1. Results in Figure 6A showed that the pairwise correlations were enhanced after imputation in the correct direction [24]. Specifically, G2S3 showed the greatest enhancement of the regulatory relationship for both inhibitory and activatory pairs. DCA enhanced more on the activatory pairs with positive correlations, scImpute did not have comparable enhancement, and MAGIC performed worse than the raw data with again a pattern of over-smoothing. We further examined the correlation enhancement of each method by plotting a set of representative inhibitory and activatory gene pairs. The gene pair *Sfpi*1 and *Gata*1 were used as an example of a mutually inhibitory relationship (Figure 6B). For this pair, scImpute did not improve the correlations. MAGIC and DCA tended to over-impute to the extent that only one gene was expressed in the same cell after imputation. This is against the observation from the raw data and previous literatures that the higher expression of one gene, the lower, rather than completely shutting off, the expression of the other. The imputed data by G2S3 showed a negatively correlated curve where the expression level of one gene decreased with the increase of the other. Another example with an activatory pair was plotted using *Zfpm*1 and *Gata*1 (Figure 6C). The positive correlation was reasonably enhanced by G2S3. For MAGIC, the plot of *Zfpm*1 and *Gata*1 formed a nearly straight diagonal line, suggesting that the imputed data was over-smoothed such that the cell-level biological variation was attenuated.

**Figure 6.**
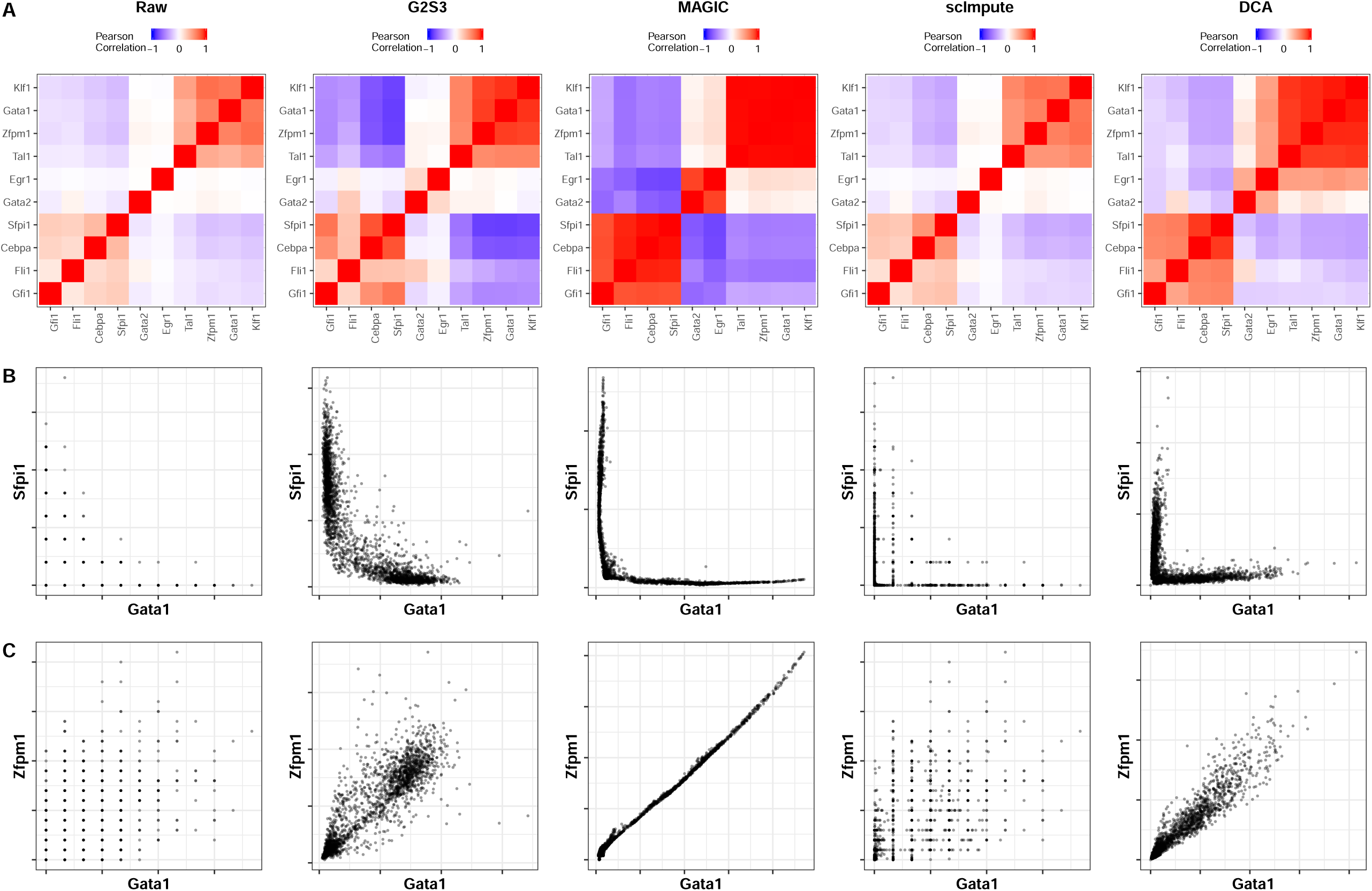
Performance of imputation methods in recovering gene regulatory relationship in the Paul dataset. **A**) heatmaps of pairwise correlations between well-known blood regulators, **B**) expression patterns of mutually inhibitory pair (*Sfpi*1 vs. *Gata*1), and **C**) expression patterns of mutually activatory pair (*Zfpm*1 vs. *Gata*1).

### Computation time

While SAVER has comparable performance to G2S3 in some datasets, G2S3 is computationally more efficient than SAVER. As both methods are based on gene networks, the computation time is expected to increase with the number of genes to be imputed. This makes gene network-based methods more suitable for scRNA-seq datasets with tens or even hundreds of thousands of cells than those based on cell similarity. In real data analysis, G2S3 was on average about six times faster than SAVER. For example, G2S3 took 11.25 hr to impute 22,934 genes from 1,529 cells in the Petropoulos dataset using an 8-core, 100-GB RAM, Intel Xeon 2.6 GHz CPU machine, whereas SAVER was unable to finish imputation within 48 hr even with the option of do.fast using the same computer. On the other hand, the computation time of the imputation methods that borrow information from similar cells increases dramatically with the number of cells in the data. As demonstrated in a previous study, scImpute and VIPER were unable to scale beyond 10K cells within 24 hr [16].

## Discussion

We have developed a novel method G2S3 to impute dropouts in scRNA-seq data. G2S3 learns a sparse and smooth signals of gene graph from scRNA-seq data and then borrows information from nearby genes in the graph for imputation. We evaluated and compared the performance of G2S3 and five existing imputation methods in terms of recovering expression levels, restoring cell subtype separation, improving differential expression analysis, and restoring gene regulatory relationships using seven scRNA-seq datasets. The results demonstrated that G2S3 achieved superior performance in all four aspects compared to the other methods, especially for genes with relatively low expression. Furthermore, G2S3 is the most computationally efficient for large-scale scRNA-seq data imputation.

Unlike the imputation methods that borrow information across similar cells, G2S3 harnesses the structural relationship among genes obtained through graph signal processing to perform imputation. Using the seven real datasets, we showed that methods relying on cell similarity tend to remove biological variation among cells and intensify subject-level batch effects. In contrast, G2S3 enhances cell subtype separation and thus relatively reduces the variations in cells from the same cell type and subject. The down-sampling and differential expression analysis results showed that G2S3 outperformed the other methods, especially for the mildly expressed genes. Of note, imputation methods such as SAVER, scImpute, and VIPER, used parametric models for gene expression. However, as the noise distribution varies across platforms for single-cell transcriptomics, the parametric model assumptions may be violated, particularly for new technologies. Graph signal processing extracts signals from data by optimizing a smoothness regulated objective function, thus is in principle less sensitive to the noise distribution.

Despite the advantages of G2S3 over the other imputation methods shown in this article, G2S3 has a number of limitations that can be improved upon. First, G2S3 uses a lazy random walk on the gene graph to recover dropout events, i.e., a weighted average of the observed expression and the predicted expression from neighboring genes. The weights currently depend only on between gene similarity which can be improved to reflect the reliability of observed read counts, the cell library size, and the dispersion of gene expression, similar to the weights used in SAVER. Second, G2S3 does not consider the dropout probability and therefore imputes all values at once. This can be improved by calculating the probability of being a dropout for each observed read count and only apply imputation on those with a high dropout probability. Finally, G2S3 does not consider the potential subject effects in the data, which has been shown to be prevalent and dominant in certain cell types. One way to address this issue is to consider subject effect as “batch” effect and remove it using batch effect removal tools. This is effective only when there are no other effects of interest, for example, disease effect, because they will also be removed with “batch” effect. When there are other effects that confound with subject effect and are the interest of study, G2S3 can be improved to consider subject effect and disease effect at the same time in imputation.

## Conclusions

In this study, we developed G2S3, an imputation method that applies graph signal processing to extract gene graph structure from scRNA-seq data and recover true expression levels by borrowing information from adjacent genes in the gene graph. G2S3 was shown to be an effective tool to improve the quality of single-cell transcriptomic data. Moreover, G2S3 is computationally efficient for imputation in large-scale scRNA-seq datasets.

## Methods

### Imputation model and optimization

To borrow information from similar genes for data imputation, G2S3 first builds a sparse graph representation of gene network under the assumption that expression levels change smoothly between closely connected genes. Let *X* = [*x*_1_, *x*_2_, …, *x*_*m*_] ∈ ℝ^*n*×*m*^ denote the observed transcript counts of *m* genes in *n* cells, where the column *x*_*j*_ ∈ ℝ^*n*^ represents the expression vector of gene *j*, for *j* = 1, …, *m*. We regard each gene *j* as a vertex *V*_*j*_ in a weighted gene graph *G* = (*V, E*), in which the edge between genes *j* and *k* is associated with a weight *W*_*jk*_.

The gene graph is then determined by the weighted adjacency matrix 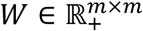. G2S3 searches for a valid adjacency matrix *W* from the space

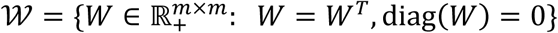

that is optimal under the assumption of smoothness and sparsity on the graph. To achieve this, we use the objective function adapted from Kalofolias’s model [30]:

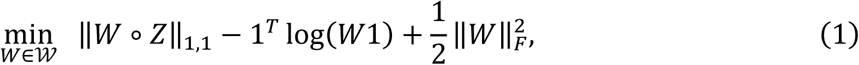

where 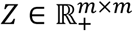 is the pairwise Euclidean distance matrix of genes, defined as *Z_jk_* = ‖*x_j_* − *x_k_*‖^2^, ‖⋅‖_1,1_ is the elementwise L-1 norm, ∘ is the Hadamard product, and ‖⋅‖_*F*_ is the Frobenius norm. The first term in Eq. (1) is equivalent to 2 tr(*X*^*T*^*X*) that quantifies how smooth the signals are on the graph, where *L* is the graph Laplacian. This term penalizes edges between distant genes, so it prefers to put a sparse set of edges between the nodes with a small distance in *Z*. The second term in Eq. (1) represents the node degree which requires the degree of each gene to be positive to improve the overall connectivity of the gene graph. The third term in Eq. (1) controls sparsity to penalize the formation of large edges between genes.

The optimization of Eq. (1) can be solved via primal dual techniques [31]. We rewrite Eq. (1) as

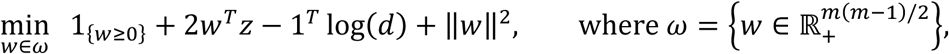

where *w* and *z* are vector forms of *W* and *Z*, respectively, *d* = *Kw* ∈ ℝ^*m*^ and *K* is the linear operator that satisfies *W*1 = *Kw*. After obtaining the optimal *W*, a lazy random walk matrix can be constructed on the graph:

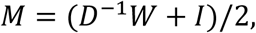

where *D* is an *m*-dimensional diagonal matrix with *D*_*jj*_ = ∑_*j*_ *W*_*jk*_, the degree of the *j*-th gene, and *I* is the identity matrix.

The imputed count matrix *X*_imputed_ is then obtained by taking one step of a random walk on the graph which can be written as

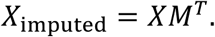

Similar to other diffusion-based methods, G2S3 spreads out counts while keeping the sum constant in the random walk step. This results in the average value of non-zero matrix entry decreasing after imputation. To match the observed expression at the gene level, we rescale the values in *X*_imputed_ so that the mean expression of each gene in the imputed data matches that of the observed data. The pseudo-code for G2S3 is given in Algorithm 1.

#### Algorithm 1: Pseudo-code G2S3

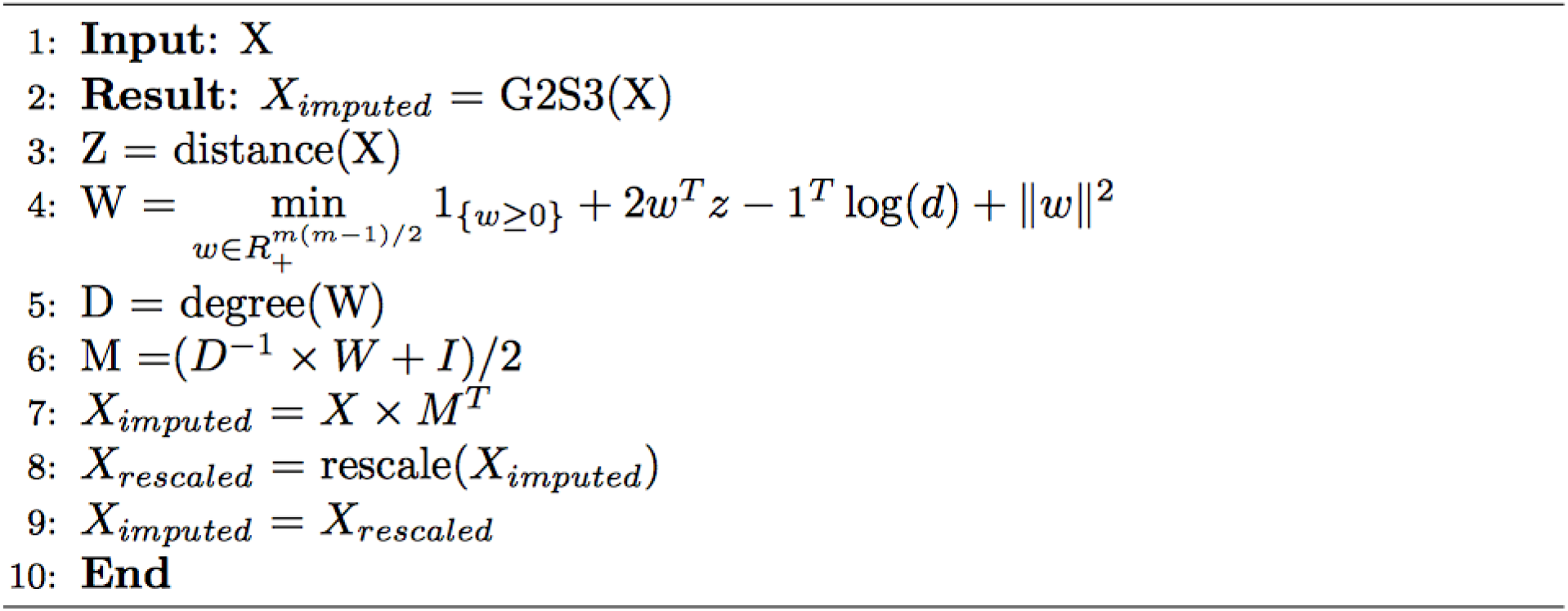

### Real datasets

We evaluated and compared the performance of G2S3 and five existing imputation methods using datasets from seven scRNA-seq studies. Among them, four datasets were generated using the UMI techniques and three were generated by non-UMI-based techniques.

**Reyfman** refers to the scRNA-seq dataset of human lung tissue from healthy transplant donors in Reyfman et al. [18]. The raw data include 33,694 genes and 5,437 cells. To generate the reference dataset, we selected cells with a total number of UMIs greater than 10,000 and genes that have nonzero expression in more than 20% of cells. This ended up with 3,918 genes and 2,457 cells.

**PBMC** refers to human peripheral blood mononuclear cells from a healthy donor stained with TotalSeq-B antibodies generated by the high-throughput droplet-based system [19]. This dataset was downloaded from 10x Genomics website (https://support.10xgenomics.com/single-cell-gene-expression/datasets). The raw data include 33,538 genes and 7,865 cells. To generate the reference dataset, we selected cells with a total number of UMIs greater than 5,000 and genes that have nonzero expression in more than 20% of cells. This ended up with 2,308 genes and 2,081 cells.

**Zeisel** refers to the scRNA-seq dataset of mouse cortex and hippocampus in Zeisel et al. [20]. The raw data include 19,972 genes and 3,005 cells. To generate the reference dataset, we selected cells with a total number of UMIs greater than 10,000 and genes that have nonzero expression in more than 40% of cells. This ended up with 3,529 genes and 1,800 cells.

**Chu** refers to the dataset investigating the separation of cell subpopulations in Chu et al. [21]. It measured gene expression of 1,018 cells including undifferentiated H1 and H9 human ES cell lines and the H1-derived progenitors. The cells were annotated with seven cell subtypes: neuronal progenitor cells (NP), definitive endoderm cells (DE), endothelial cells (EC), trophoblast-like cells (TB), human foreskin fibroblasts (HF), and undifferentiated H1 and H9 human ES cells. We performed preliminary filtering to remove genes expressed in less than 10% of cells, which resulted in 13,829 genes.

**Petropoulos** refers to the dataset studying cell lineage in human embryo development in Petropoulos et al. [22]. It measured expression profiles of 26,178 genes in 1,529 cells from 88 human embryos. Cells were labeled as E3-E7 representing their embryonic day. We performed preliminary filtering to remove genes expressed in less than 5 cells and cells with less than 200 expressed genes. After the filtering, we ended up with 22,934 genes and 1,529 cells.

**Segerstolpe** refers to the dataset in Segerstolpe et al. [23]. The original dataset includes 26,179 genes and 3,514 cells from whole islets of ten subjects, among which four are T2D patients and six are healthy donors. Samples from the four T2D patients and three healthy donors were also processed for bulk RNA-seq. We used the seven subjects with both scRNA-seq and bulk RNA-seq data to evaluate the method performance in improving differential expression analysis. We performed preliminary filtering to remove genes expressed in less than 20% of cells, which resulted in 9,010 genes.

**Paul** refers to the dataset from a study on the transcriptional differentiation landscape of myeloid progenitors [24]. This dataset includes 3,451 informative genes and 2,730 cells. We used this dataset to evaluate the performance of imputation methods in restoring gene regulatory relationships between well-known regulators.

### Performance evaluation

#### Expression data recovery

We first compared the method performance in recovering true expression levels using down-sampled datasets. Down-sampling was performed on three independent UMI-based scRNA-seq datasets (Reyfman, PBMC, and Zeisel) to generate benchmarking datasets in a similar framework to previous studies [13,16]. In each dataset, we selected a subset of genes and cells with high expression to be used as the reference dataset and treated them as the true expression. Details on the thresholds chosen to generate the reference datasets are described in the “Real datasets” section. However, unlike previous studies that simulated down-sampled datasets from models with certain distributional assumptions [13] which may incur modeling bias, we performed random binary masking of UMIs in the reference datasets to mimic the inefficient capturing of transcripts in dropout events. The binary masking process masked out each UMI independently with a given probability. In each reference dataset, we randomly masked out 80% of UMIs to create the down-sampled dataset.

All imputation methods were applied to each down-sampled dataset to generate imputed data separately. Because imputation methods such as SAVER and MAGIC output the normalized library size values, we performed library size normalization on all imputation methods. We calculated the gene-wise Pearson correlation and cell-wise Spearman correlation between the reference data and the imputed data generated by each imputation method. The correlations were also calculated between the reference data and the observed data without imputation to provide a baseline for comparison. To investigate whether the performance depends on the true expression level, we stratified genes into three categories: widely, mildly, and rarely expressed genes, based on the proportion of cells expressing genes in the down-sampled datasets. Specifically, widely expressed genes are those with non-zero expression in more than 80% of cells, rarely expressed genes are those with non-zero expression in less than 30% of cells, and mildly expressed genes are those that lie in between. The gene-wise and cell-wise correlations in each stratum were used to demonstrate the impact of expression level on the performance of imputation methods.

#### Restoration of cell subtype separation

We applied the six imputation methods to the Chu dataset to evaluate their performance in separating different cell types. Methods that took more than 24 hr to finish imputation were excluded. A good imputation method is expected to stabilize within cell-subtype variation (intra-subtype distance) while maintaining between cell-subtype variation (inter-subtype distance). Principal component analysis was conducted on the raw and imputed data for dimension reduction. We calculated the inter-subtype distance as the Euclidian distance between cells from different cell types, and the intra-subtype distance as the distance between cells of the same cell type, using the top *K* PCs of the data, for *K* = 1, …, 10. The ratio of the average inter-subtype distance to the average intra-subtype distance was used to quantify the performance. The higher this ratio is, the better performance the method has. We also calculated the silhouette coefficient, a composite index reflecting both the compactness and separation of different cell types, using the top *K* PCs. The silhouette coefficient ranges from −1 to 1 where a higher value indicates a better matching with the cell subtypes and a value close to zero indicates random clustering [32]. To demonstrate the comparison using cell clustering results, we visualized the raw and imputed data using the top three PCs and colored cells by the cell subtype labels. To demonstrate cell subtype separation based on cell subtype marker genes, we further displayed DE and H1/H9 cells using their marker genes [21]: *GATA*6, a marker gene of DE cells, and *NANOG*, a marker gene of H1/H9 cells.

We then assessed the performance of G2S3 and MAGIC in restoring the separation of cells from human preimplantation embryos of different embryonic days in the Petropoulos dataset. Since cell developmental time is the primary segregating factor for cells [21], a good imputation method is expected to restore the separation of cells from different embryonic days instead of from different subjects. Cells were clustered using the Louvain algorithm [25] in the observed and imputed data by G2S3 and MAGIC. The performance was evaluated based on data visualization using t-SNE plots with cells colored by the detected cell clusters, the embryonic days, and the individual origins.

#### Improvement in differential expression analysis

To assess the performance in improving the identification of differentially expressed genes, we compared T2D patients to healthy donors using both imputed scRNA-seq and bulk RNA-seq data from the Segerstolpe dataset. Differential analysis in the bulk RNA-seq data was performed using t-test and Wilcoxon rank sum test, and the differential analysis in the scRNA-seq data was performed using the Seurat R package (version 3.0) with a default threshold on genes with at least 0.25 log fold change. We treated the differentially expressed genes identified from the bulk RNA-seq data as ground truth. The predictive power of differentially expressed genes identified in the raw and imputed scRNA-seq data on the ground truth was measured by the area under an ROC curve. Genes were stratified into two groups: highly expressed (expressed in more than 30% of cells) and lowly expressed (expressed in less than 30% cells), to demonstrate the impact of expression level on the method performance.

#### Gene regulatory relationship restoration

We finally evaluated the method performance by investigating the enhancement in the discovery of gene regulatory relationships using the Paul dataset. This was achieved by assessing the pairwise correlations among a set of regulators with known inhibitory and activatory relationships in blood development [15]. The estimated pairwise correlations between genes using the raw unimputed and imputed data by each method were compared for performance evaluation. Methods that took more than 24 hr to finish imputation were excluded from the results.

## Abbreviations

scRNA-seq: Single-cell RNA sequencing
UMI: Unique Molecular Identifier
PBMC: Peripheral Blood Mononuclear Cell
ES: Embryonic Stem
T2D: Type II Diabetes
PC: Principal Component
t-SNE: t-Distributed Stochastic Neighbor Embedding
ROC: Receiver Operating Characteristic
AUC: Area Under the Curve

## Declarations

### Availability of data and materials

G2S3 is an open-source MATLAB package that is freely available on GitHub https://github.com/ZWang-Lab/G2S3 under the MIT license. The detailed list of data sets used in the study is described in the “Real datasets” section. The code to reproduce all the analyses presented in the paper are available on GitHub https://github.com/ZWang-Lab/G2S3_paper2020.

### Authors’ contributions

WW, XY, and ZW conceived the idea, developed the method, and designed the study. WW implemented the software and performed the analyses. QD and YL contributed to data collection. WW, XY, and ZW wrote the manuscript. XY and ZW supervised the research. All authors read and approved the final manuscript.

### Ethics approval and consent to participate

No ethical approval was required for this study. All public datasets used in the paper were generated by other organizations that have obtained ethical approval.

## Acknowledgements

The authors would like to thank Drs. Smita Krishnaswamy and Hongyu Zhao for their valuable suggestions and comments.

## Consent for publication

Not applicable

## References

1. Grün D, Lyubimova A, Kester L, Wiebrands K, Basak O, Sasaki N, et al. Single-cell messenger RNA sequencing reveals rare intestinal cell types. Nature. 2015;525:251–5.

2. Mahata B, Zhang X, Kolodziejczyk AA, Proserpio V, Haim-Vilmovsky L, Taylor AE, et al. Single-Cell RNA Sequencing Reveals T Helper Cells Synthesizing Steroids De Novo to Contribute to Immune Homeostasis. Cell Reports. 2014;7:1130–42.

3. Usoskin D, Furlan A, Islam S, Abdo H, Lönnerberg P, Lou D, et al. Unbiased classification of sensory neuron types by large-scale single-cell RNA sequencing. Nat Neurosci. 2015;18:145–53.

4. Frishberg A, Peshes-Yaloz N, Cohn O, Rosentul D, Steuerman Y, Valadarsky L, et al. Cell composition analysis of bulk genomics using single-cell data. Nat Methods. 2019;16:327–32.

5. Wu YE, Pan L, Zuo Y, Li X, Hong W. Detecting Activated Cell Populations Using Single-Cell RNA-Seq. Neuron. 2017;96:313-329.e6.

6. Yuan G-C, Cai L, Elowitz M, Enver T, Fan G, Guo G, et al. Challenges and emerging directions in single-cell analysis. Genome Biology. 2017;18:84.

7. Shalek AK, Benson M. Single-cell analyses to tailor treatments. Science Translational Medicine [Internet]. 2017 [cited 2019 Nov 7];9. Available from: https://stm.sciencemag.org/content/9/408/eaan4730

8. Kharchenko PV, Silberstein L, Scadden DT. Bayesian approach to single-cell differential expression analysis. Nat Methods. 2014;11:740–2.

9. van Dijk D, Sharma R, Nainys J, Yim K, Kathail P, Carr AJ, et al. Recovering Gene Interactions from Single-Cell Data Using Data Diffusion. Cell. 2018;174:716–729.e27.

10. Li WV, Li JJ. An accurate and robust imputation method scImpute for single-cell RNA-seq data. Nature Communications. 2018;9:1–9.

11. Gong W, Kwak I-Y, Pota P, Koyano-Nakagawa N, Garry DJ. DrImpute: imputing dropout events in single cell RNA sequencing data. BMC Bioinformatics. BioMed Central; 2018;19:220.

12. Chen M, Zhou X. VIPER: variability-preserving imputation for accurate gene expression recovery in single-cell RNA sequencing studies. Genome Biology. BioMed Central; 2018;19:196.

13. Huang M, Wang J, Torre E, Dueck H, Shaffer S, Bonasio R, et al. SAVER: Gene expression recovery for single-cell RNA sequencing. Nature Methods. Springer US; 2018;15:539–42.

14. Talwar D, Mongia A, Sengupta D, Majumdar A. AutoImpute: Autoencoder based imputation of single-cell RNA-seq data. Scientific Reports. 2018;8:16329.

15. Eraslan G, Simon LM, Mircea M, Mueller NS, Theis FJ. Single-cell RNA-seq denoising using a deep count autoencoder. Nature Communications. 2019;10:390.

16. Arisdakessian C, Poirion O, Yunits B, Zhu X, Garmire LX. DeepImpute: an accurate, fast and scalable deep neural network method to impute single-cell RNA-Seq data. bioRxiv. 2018;353607.

17. Andrews TS, Hemberg M. False signals induced by single-cell imputation. F1000Res [Internet]. 2019 [cited 2019 Sep 30];7. Available from: https://www.ncbi.nlm.nih.gov/pmc/articles/PMC6415334/

18. Reyfman PA, Walter JM, Joshi N, Anekalla KR, McQuattie-Pimentel AC, Chiu S, et al. Single-Cell Transcriptomic Analysis of Human Lung Provides Insights into the Pathobiology of Pulmonary Fibrosis. American Journal of Respiratory and Critical Care Medicine. 2018;rccm.201712-2410OC.

19. Zheng GXY, Terry JM, Belgrader P, Ryvkin P, Bent ZW, Wilson R, et al. Massively parallel digital transcriptional profiling of single cells. Nat Commun. 2017;8:1–12.

20. Zeisel A, Munoz-Manchado AB, Codeluppi S, Lonnerberg P, La Manno G, Jureus A, et al. Cell types in the mouse cortex and hippocampus revealed by single-cell RNA-seq. Science. 2015;347:1138–42.

21. Chu L-F, Leng N, Zhang J, Hou Z, Mamott D, Vereide DT, et al. Single-cell RNA-seq reveals novel regulators of human embryonic stem cell differentiation to definitive endoderm. Genome Biology. 2016;17:173.

22. Petropoulos S, Edsgärd D, Reinius B, Deng Q, Panula SP, Codeluppi S, et al. Single-Cell RNA-Seq Reveals Lineage and X Chromosome Dynamics in Human Preimplantation Embryos. Cell. 2016;165:1012–1026.

23. Segerstolpe Å, Palasantza A, Eliasson P, Andersson E-M, Andréasson A-C, Sun X, et al. Single-Cell Transcriptome Profiling of Human Pancreatic Islets in Health and Type 2 Diabetes. Cell Metab. 2016;24:593–607.

24. Paul F, Arkin Y, Giladi A, Jaitin DA, Kenigsberg E, Keren-Shaul H, et al. Transcriptional Heterogeneity and Lineage Commitment in Myeloid Progenitors. Cell. 2015;163:1663–77.

25. Blondel VD, Guillaume J-L, Lambiotte R, Lefebvre E. Fast unfolding of communities in large networks. J Stat Mech. 2008;2008:P10008.

26. Levine JH, Simonds EF, Bendall SC, Davis KL, Amir ED, Tadmor MD, et al. Data-Driven Phenotypic Dissection of AML Reveals Progenitor-like Cells that Correlate with Prognosis. Cell. 2015;162:184–97.

27. Krumsiek J, Marr C, Schroeder T, Theis FJ. Hierarchical Differentiation of Myeloid Progenitors Is Encoded in the Transcription Factor Network. Pesce M, editor. PLoS ONE. 2011;6:e22649.

28. Rekhtman N, Radparvar F, Evans T, Skoultchi AI. Direct interaction of hematopoietic transcription factors PU.1 and GATA-1: functional antagonism in erythroid cells. Genes Dev. 1999;13:1398–411.

29. Iwasaki H, Mizuno S, Wells RA, Cantor AB, Watanabe S, Akashi K. GATA-1 Converts Lymphoid and Myelomonocytic Progenitors into the Megakaryocyte/Erythrocyte Lineages. Immunity. 2003;19:451–62.

30. Kalofolias V. How to Learn a Graph from Smooth Signals. Artificial Intelligence and Statistics [Internet]. 2016 [cited 2020 Mar 21]. p. 920–9. Available from: http://proceedings.mlr.press/v51/kalofolias16.html

31. Komodakis N, Pesquet J-C. Playing with Duality: An Overview of Recent Primal-Dual Approaches for Solving Large-Scale Optimization Problems. 2014;

32. Rousseeuw PJ. Silhouettes: A graphical aid to the interpretation and validation of cluster analysis. Journal of Computational and Applied Mathematics. 1987;20:53–65.

