## Supplemental figures for "G2S3: a gene graph-based imputation method for single-cell RNA sequencing data"

by

Weimiao Wu, Qile Dai, Yunqing Liu, Xiting Yan, Zuoheng Wang

### Supplemental Figures

**Figure S1:** Performance of imputation methods measured by correlation with reference data in three down-sampled datasets, on the gene level (top) and cell level (bottom). Genes are stratified into three groups: widely (>80%, left), mildly (30%-80%, middle), and rarely (<30%, right) expressed.

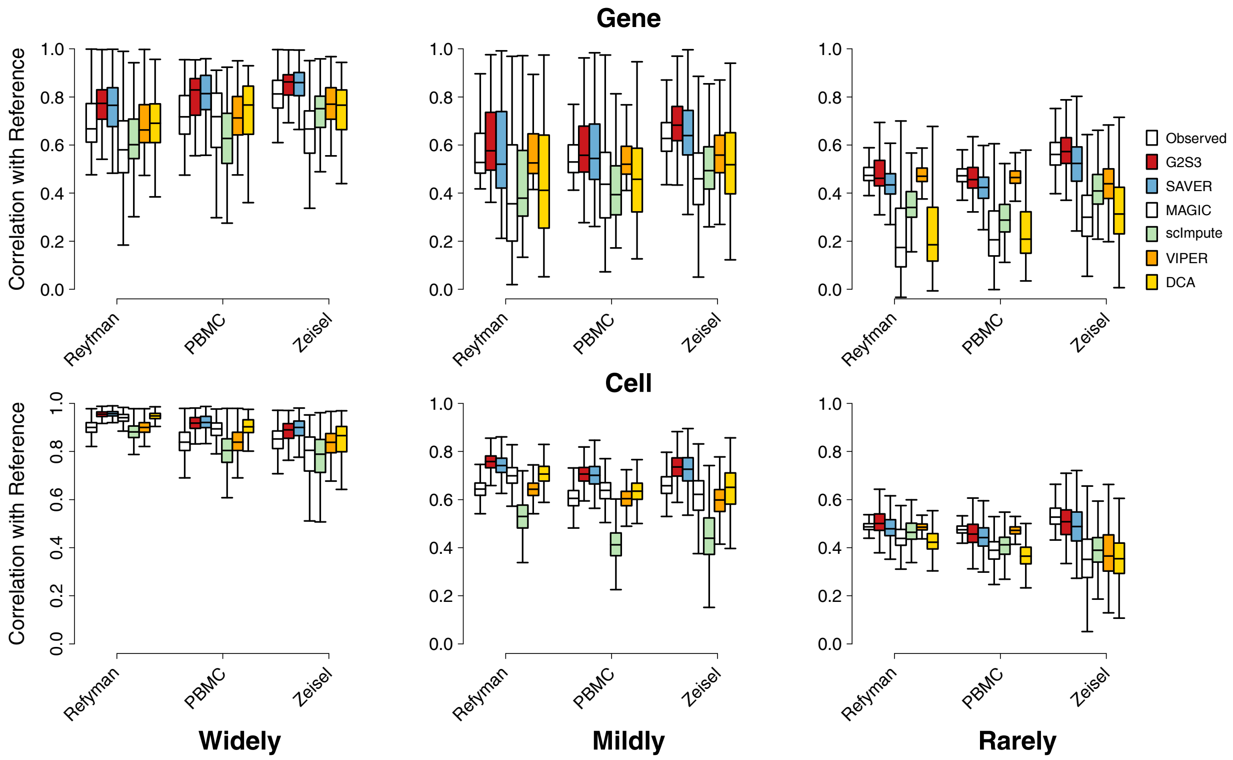

**Figure S2:** PC plots (PC1 vs. PC2) of the raw unimputed and imputed data by each method in the Chu dataset. Cells are colored by the cell subtype labels.

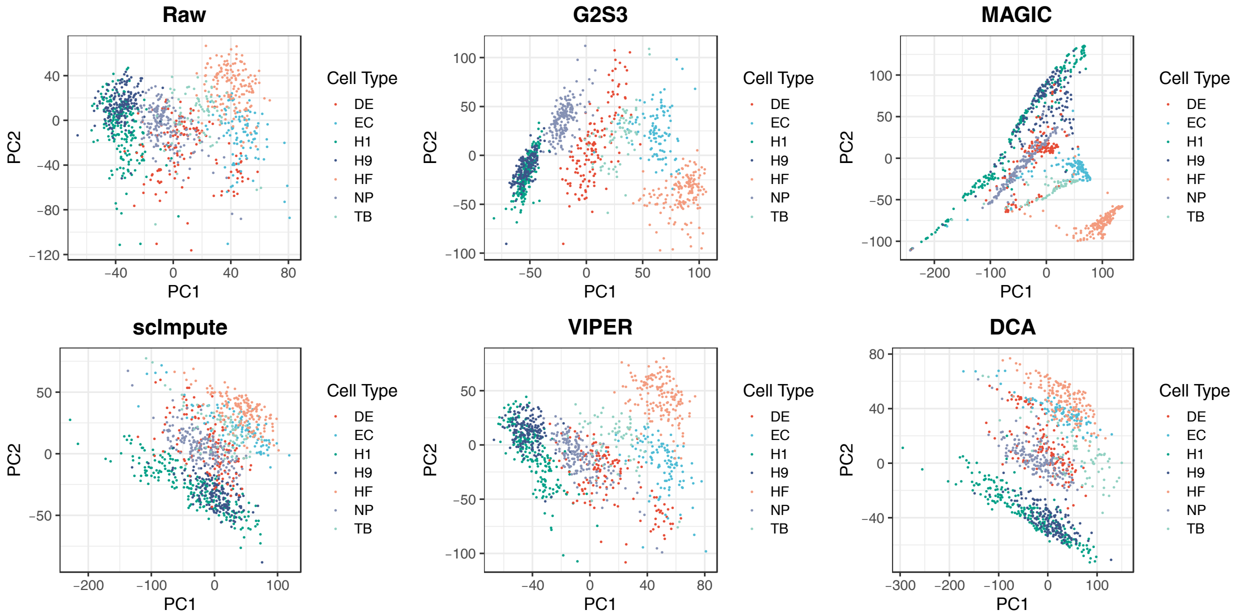

**Figure S3:** Scatter plot of expression on marker genes for DE cells (*GATA6*) and H1/H9 cells (*NANOG*) in the Chu dataset. Cells are colored by the cell subtype labels.

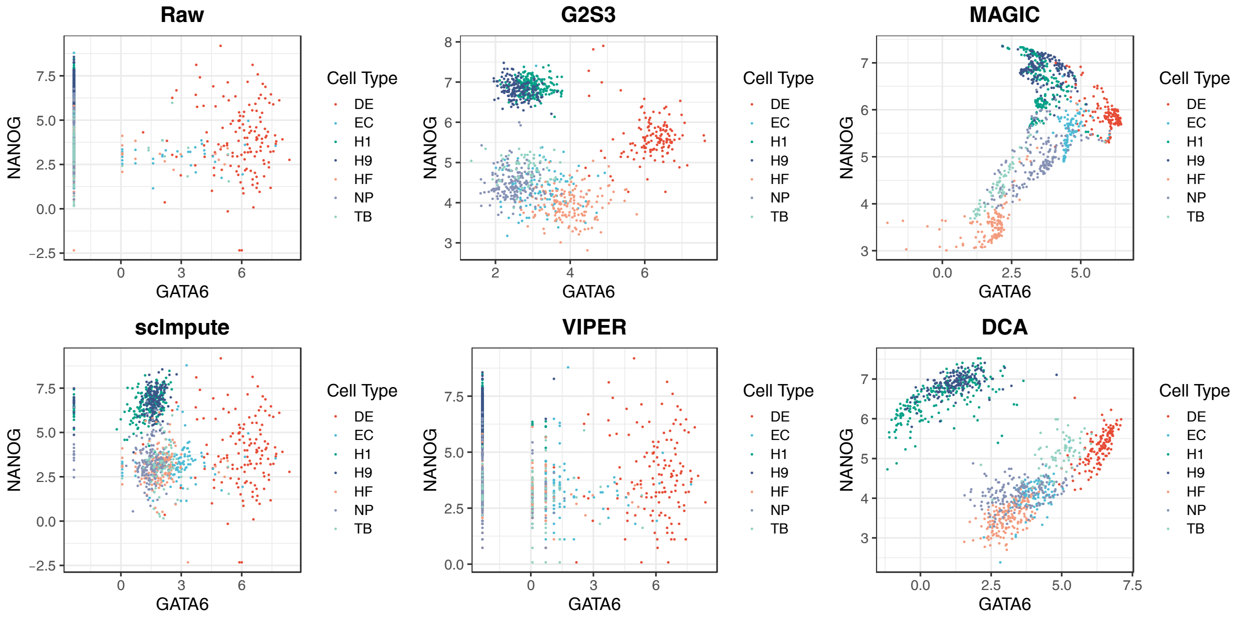
